# Loss of coordinated expression between ribosomal and mitochondrial genes revealed by comprehensive characterization of a large family with a rare mendelian disorder

**DOI:** 10.1101/2020.10.22.350884

**Authors:** Brendan Panici, Hosei Nakajima, Colleen M. Carlston, Hakan Ozadam, Can Cenik, Elif Sarinay Cenik

**Affiliations:** Department of Molecular Biosciences, University of Texas at Austin, Austin, USA; School of Medicine, University of California, San Francisco, San Francisco, CA

## Abstract

Non-canonical intronic variants are a poorly characterized yet highly prevalent class of alterations associated with Mendelian disorders. Here, we report the first RNA expression and splicing analysis from a family whose members carry a non-canonical splice variant in an intron of *RPL11* (c.396+3A>G). This mutation is causative for Diamond Blackfan Anemia (DBA) in this family despite incomplete penetrance and variable expressivity. Our analyses revealed a complex pattern of disruptions with many novel junctions of *RPL11*. These include an *RPL11* transcript that is translated with a late stop codon in the 3’ untranslated region (3’UTR) of the main isoform. We observed that *RPL11* transcript abundance is comparable among carriers regardless of symptom severity. Interestingly, both the small and large ribosomal subunit transcripts were significantly overexpressed in individuals with a history of anemia in addition to congenital abnormalities. Finally, we discovered that coordinated expression between mitochondrial components and *RPL11* was lost in all carriers, which may lead to variable expressivity. Overall, this study highlights the importance of RNA splicing and expression analyses in families for molecular characterization of Mendelian diseases.

## INTRODUCTION

Many genetic diseases are incompletely penetrant and exhibit variable expressivity. The nature of the causative genetic mutation, epistastic interactions, and environmental factors can accentuate, mitigate, or sometimes completely mask a potentially pathogenic genetic variant. Diamond Blackfan Anemia (DBA) is an example of a monoallelic, genetic disease exhibiting incomplete penetrance and variable expressivity (reviewed in [1]). DBA can lead to developmental and hematological defects that often manifest in the first year of infancy and is caused by haploinsufficiency in the protein synthesis machinery due to loss of function or missense mutations. DBA mutations have been identified in a ribosome maturation factor, *TSR2*, two transcriptional regulators, *GATA1*, and *MYSM1*, and ∼19 ribosomal protein genes including *RPL11* [2–5].

*RPL11* mutations in DBA patients are associated with a spectrum of phenotypes. Compared to other DBA genes such as *RPS19*, individuals with *RPL11* mutations have an increased incidence of cleft palate and thumb anomalies [6,7]. Loss of one functional copy of *RPL11* leads to impaired red blood cell development, reduced P53, increased c-MYC in mouse [8], and in contradiction increased P53 and apoptosis in zebrafish [9]. *RPL11* mutations are relatively common in DBA patients affecting ∼ 7%. In comparison, *RPS19* mutations are the most prevalent cause (∼30-38% of patients) [5].

While initial characterizations of DBA patients focused on nonsense and missense coding sequence mutations in ribosomal proteins, recent evidence indicates that extended splice site variants are under-appreciated and highly prevalent [5]. Splicing variants may disrupt a splicing site, activate a cryptic/alternative splice isoform, or introduce novel junctions with adverse consequences for the open reading frame of the corresponding gene [10]. Genome and exome sequencing in patients revealed many potentially pathogenic intronic variants that may alter splicing. For example, a rare, synonymous variant in *GFI1B*–a transcription regulator expressed in hematopoietic lineage– is responsible for variation in blood traits across humans [11]. Despite the high prevalence of splice site variants in DBA patients (∼18.5% [5]), no previous study has characterized the RNA expression and splicing changes in patient samples harboring such splice variants.

Here, we report the first detailed RNA expression and splicing analysis in a large family with a noncanonical intronic splice variant of the ribosomal protein gene, *RPL11*. Eight family members spanning four generations harbor the particular non-canonical *RPL11* splicing variant (c.396+3A>G). In this family, two carriers have prominent congenital symptoms (congenital heart defect & short stature) and the other carriers are either asymptomatic or have small thumbs that doesn’t require any surgical intervention [12]. This particular RPL11 splicing variant had also been observed in an individual with bilateral triphalangeal thumbs, heart defects, short stature and anemia [13].

In this study, we address the following questions: How does the pattern of *RPL11* splicing differ in carriers compare to noncarrier relatives? Is there a shared gene expression signature among *RPL11* splicing variant carriers? Do we observe any significant changes in gene expression patterns in carrier individuals with a history of anemia compared to *RPL11* variant carriers with few to no symptoms?

Our analyses revealed a complex pattern of splicing disruptions including an *RPL11* transcript that is translated with a late stop codon in the 3’ untranslated region (3’UTR) of the main isoform. *RPL11* transcript abundance is comparable among carriers regardless of symptom severity. Interestingly, both the small and large ribosomal subunit transcripts were significantly overexpressed in individuals with a history of anemia in addition to congenital abnormalities. Finally, we observed that coordinated expression between mitochondrial components and *RPL11* was lost in all carriers, which may lead to variable expressivity. Overall, this study highlights the importance of RNA splicing and expression analyses in families for molecular characterization of Mendelian diseases.

## RESULTS

### RNA sequencing analysis of a large family with a noncanonical splice variant in *RPL11*

We performed whole blood RNA expression and splicing analysis from members of a family spanning four generations that harbor an intronic splicing variant in ribosomal protein gene *RPL11* (**Figure-1**) [12]. Family members carrying the *RPL11* variant (light red circles/squares) that are not affected by anemia had small thumbs or thenar eminences that did not require surgical intervention, and one carrier of the familial variant was asymptomatic. Clinical findings in the carrier individuals with a history of anemia (dark red circle and square) are as follows: Individual IV.5 was affected with bilateral thumb defects, anemia, short stature, congenital heart defects (ASD and double superior vena cava), peno-scrotal transposition hypospadias and volvulus. Individual III.13 has cleft palate, anemia, celiac-like gastrointestinal issues and slender but not hypoplastic thumbs. At the time samples were taken for RNA extraction, both individuals had mild anemia, since that time individual IV.5’s hemoglobin continued to rise to now be within the lower range of normal.

**Figure-1:**
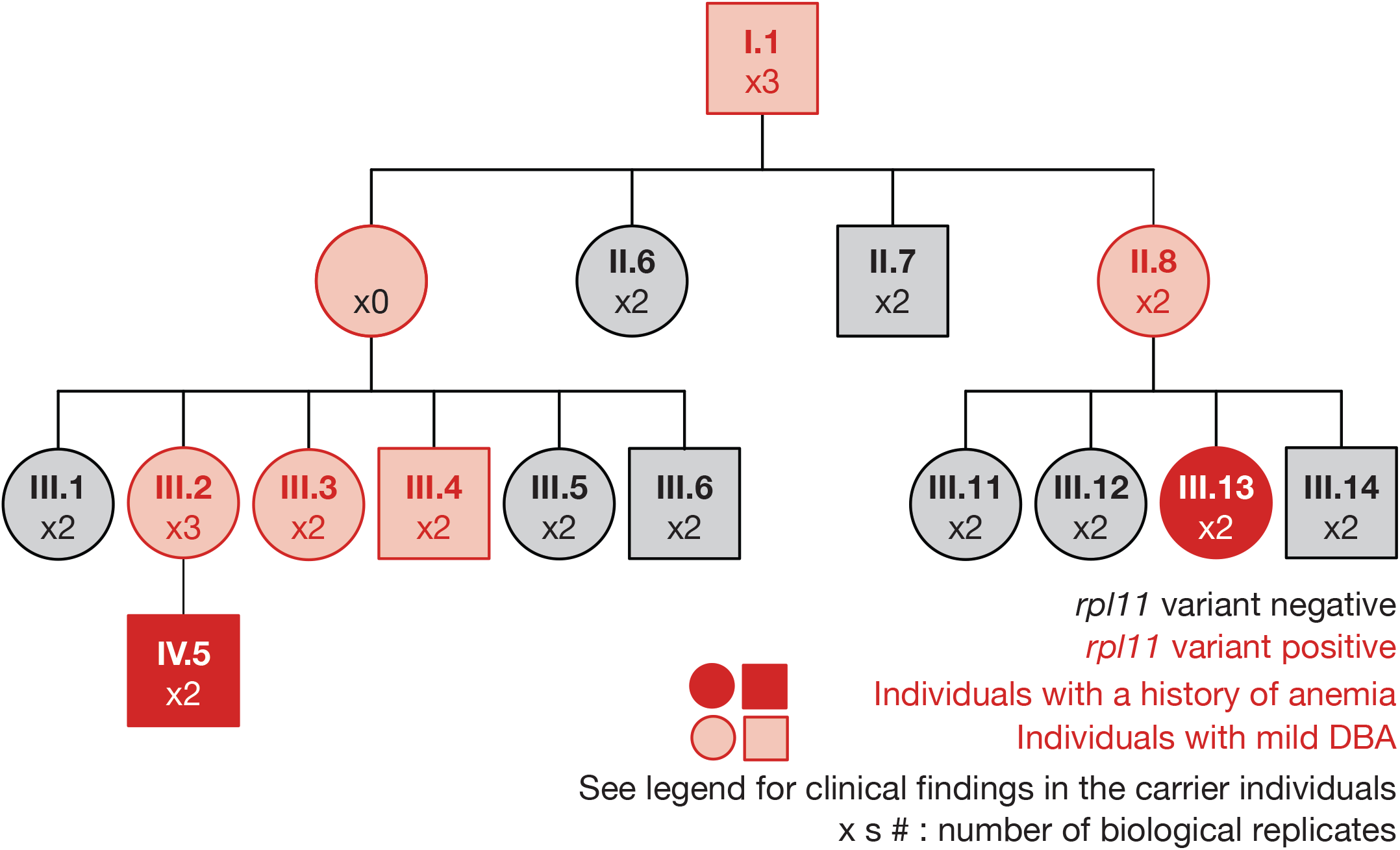
Four generation family pedigree depicts individuals who carry the pathogenic *RPL11* variant c.396+3A>G [adapted and simplified from (Carlston et al., 2017)]. Light red circles and squares represent individuals who carry the RPL11 variant. Clinical findings in the carrier individuals with a history of anemia (dark red circle and square) are as follows: Individual IV.5 was affected with bilateral thumb defects, anemia, short stature, congenital heart defects (ASD and double superior vena cava), peno-scrotal transposition hypospadias and volvulus. Individual III.13 has cleft palate, anemia, celiac-like gastrointestinal issues and slender but not hypoplastic thumbs. Other family members carrying the *RPL11* variant (light red circles/squares) had small thumbs or thenar eminences that did not require surgical intervention, and one carrier of the familial variant was asymptomatic (annotated as individuals with mild DBA).

Using our RNA sequencing data, we first confirmed the presence of the intronic variant (c.396+3A>G) for each carrier (**Supplemental Figure-1**). Given that all samples are generated from the same biological material source and family, we expected to observe global similarity in expression profiles across the samples. We identified five samples that behaved differently compared to the rest of the samples (**Supplemental Figure-2**, mean spearman correlation of outlier samples <0.9). These outlier samples predominantly had low percentage of genome mapping reads and were removed from further analyses (**Supplemental Figure-3**). The overall mean correlation coefficient was 0.97 between remaining samples (**Supplemental Figure-2**).

After sequencing depth normalization (median of ratios based on DESeq2), we didn’t observe any bias for total normalized read counts per sample (**Supplemental Figure-4A**). Principal component analyses revealed that RNA-Seq read counts from samples of the same individual tended to cluster suggesting that inter-individual variance in gene expression dominates technical variability in our data (**Supplemental Figure-4B**). Given the large number of samples, library preparation was carried out across several batches as described in **Supplemental Table-1**. Principal component analysis (PCA) suggested that batch variables do not explain a significant component of the variation observed in read counts (**Supplemental Figure-4C**). Taken together, 39 samples passed our strict quality control metrics and were used for all subsequent analyses.

### The most abundant RPL11 alternate splicing isoform results in a frame shift and a 3’UTR readthrough

Diamond Blackfan Anemia (DBA) in this family is caused by an *RPL11* 5’splice site (5’SS) variant (c.396+3A>G) (**Figure-2A, Supplementary Figure-1**). This variant is not in the canonical splice donor site, and thus we first characterized the consequent perturbation in the splicing of the RPL11 gene. Specifically, we hypothesized that differences in splicing patterns of RPL11 transcript may underlie differences in symptom among carriers.

**Figure-2:**
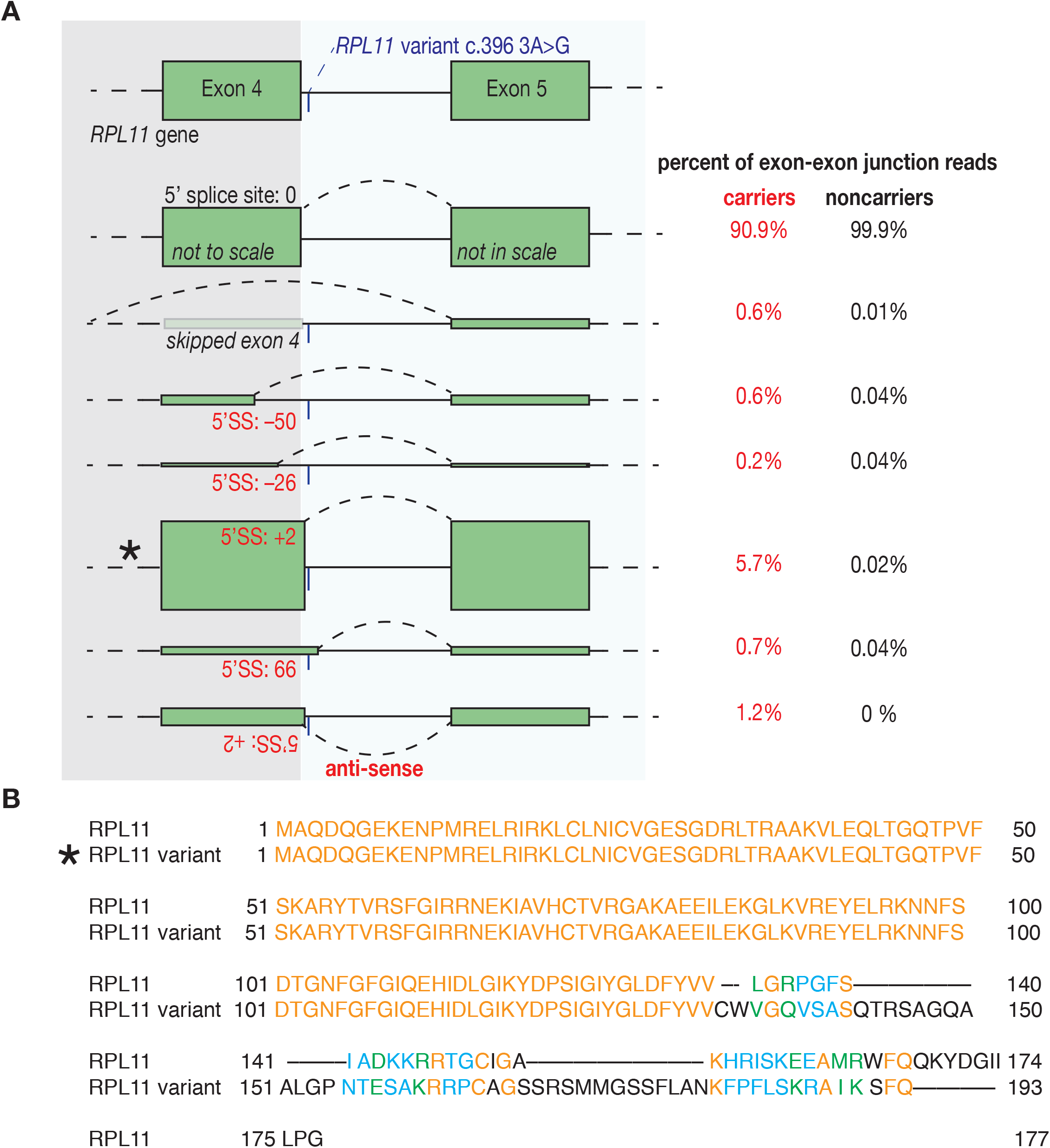
*RPL11* variant c.396+3A>G alters the splicing pattern of *RPL11*. **A** Diagram depicts exon 4–exon 5 junction RNA reads with percentages from carriers and noncarriers of the pathogenic *RPL11* variant. Left, green boxes represent exon 4 and exon 5, height of the boxes represent read abundance unless noted otherwise. Start and end positions are given in comparison to splicing isoform that produces functional canonical RPL11 protein (start, end:0). The numbers of reads mapping to each junction per individual are listed in Supplemental Table-2. The most abundant splicing variant (marked with *) results in a frame-shift and a 3’UTR ribosome readthrough. **B** Alignment of canonical RPL11 and the protein encoded by the most abundant alternate isoform was shown where amino acids are colored based on similarity.

We classified all exon junction reads proximal to the RPL11 variant with respect to the 5’splice site position (5’SS) denoting the donor site of the canonical RPL11 isoform as +0 (**Figure 2A; Supplemental Table-2**). We found that both the types of aberrant splice junctions and their relative abundances were very similar among all carrier individuals regardless of symptom severity, suggesting that the severity of symptoms is not associated with the frequency or type of aberrant splicing event.

The most abundant minor isoform in carriers has an alternative 5’ splice site at the +2 position (5.7% in carriers compared to 0.02% in noncarriers, **Figure 2A**). A previous study of samples carrying the same variant had revealed a mis-splicing event with an alternative 5’SS at the +66 position relative to the canonical 5’SS [12]. This report relied on PCR amplification of RPL11 cDNA and Sanger sequencing of a band isolated from a DNA gel thus likely missed the most abundant +2 5’SS variant. Importantly, the isoform containing the 5’SS at +2 results in a frame-shift with a stop codon in the 3’UTR of the RPL11 canonical transcript. Thus, this isoform is unlikely to be an NMD substrate due to an absence of early stop codon upstream of any intron [14]. This alternative RPL11 isoform when translated shares the first 134 amino acids with the canonical RPL11 protein (**Figure 2B**). We currently do not know whether this aberrant splice variant is translated into a stable protein as 3’UTR ribosome readthrough can lead to mRNA degradation via no-go decay (NGD; reviewed in [15] or protein instability [16], **Figure 2B**).

In addition to the most abundantly observed +2 5’SS variant, we detected ∼80 nucleotide antisense exon-4–exon-5 junction reads from the +2 5’SS (1.2% of total exon-exon junction reads). Strikingly, these reads are specifically observed in carriers but not in noncarrier individuals (**Figure 2A**). Importantly, every single sample carrying the RPL11 variant expressed the antisense exon-exon junction reads **(Supplemental Table-2)**. The alternative antisense exon-exon junctions are observed in multiple uniquely mapped reads with distinct end coordinates in ∼20 different RNA-Seq libraries from *RPL11* variant carriers, and thus are not attributable to alignment or library preparation artifacts. Furthermore, no other regions in the human genome have sequence similarity to the antisense reads. We wondered whether these junction reads could originate from a spliced antisense transcript; however, we did not observe any reads besides those mapping to the exon-exon junction on the antisense strand overlapping the RPL11 locus. Taken together, these findings reveal an intriguing antisense RNA product that is specifically produced from the RPL11 splice variant containing allele.

### Shared gene expression changes among individuals that carry the RPL11 variant c.396+3A>G

We next sought to determine differential gene expression patterns among carriers compared to noncarriers. For this purpose, we used two complementary approaches (DESeq2 and Limma-dream; see Methods). *RPL11* was among the set of differentially expressed genes with an estimated abundance of ∼%57 in carriers compared to noncarriers (DESeq2 log-fold change is 0.8, noncarriers to carriers). We hypothesized that *RPL11* levels might be different among carriers with mild symptoms vs those with a history of anemia but failed to observe any difference in *RPL11* expression. Hence, RPL11 abundance differences in carriers are unlikely to explain DBA symptom severity in these individuals.

We identified nine genes with RNA expression differences among carriers compared to noncarriers. (DESeq2 adjusted p-value < 0.01; limma-dream p-value < 0.05, **Figure-3A and B**). Interestingly, when we restricted the RNA expression comparison to only individuals with mild DBA symptoms compared to noncarriers, we identified 23 genes with RNA expression differences (DESeq2 adjusted p-value < 0.01; limma-dream p-value < 0.05, **Figure-S5A**). Genes that are differentially expressed between carriers with mild symptoms and noncarriers were surprisingly not differentially expressed in individuals with a history of anemia (**Figure-S5B; Supplemental Table-3 and Supplemental Table-4)**. This result suggests the existence of additional compensatory changes only in individuals with mild or no DBA symptoms (small thumb or asymptomatic).

**Figure-3:**
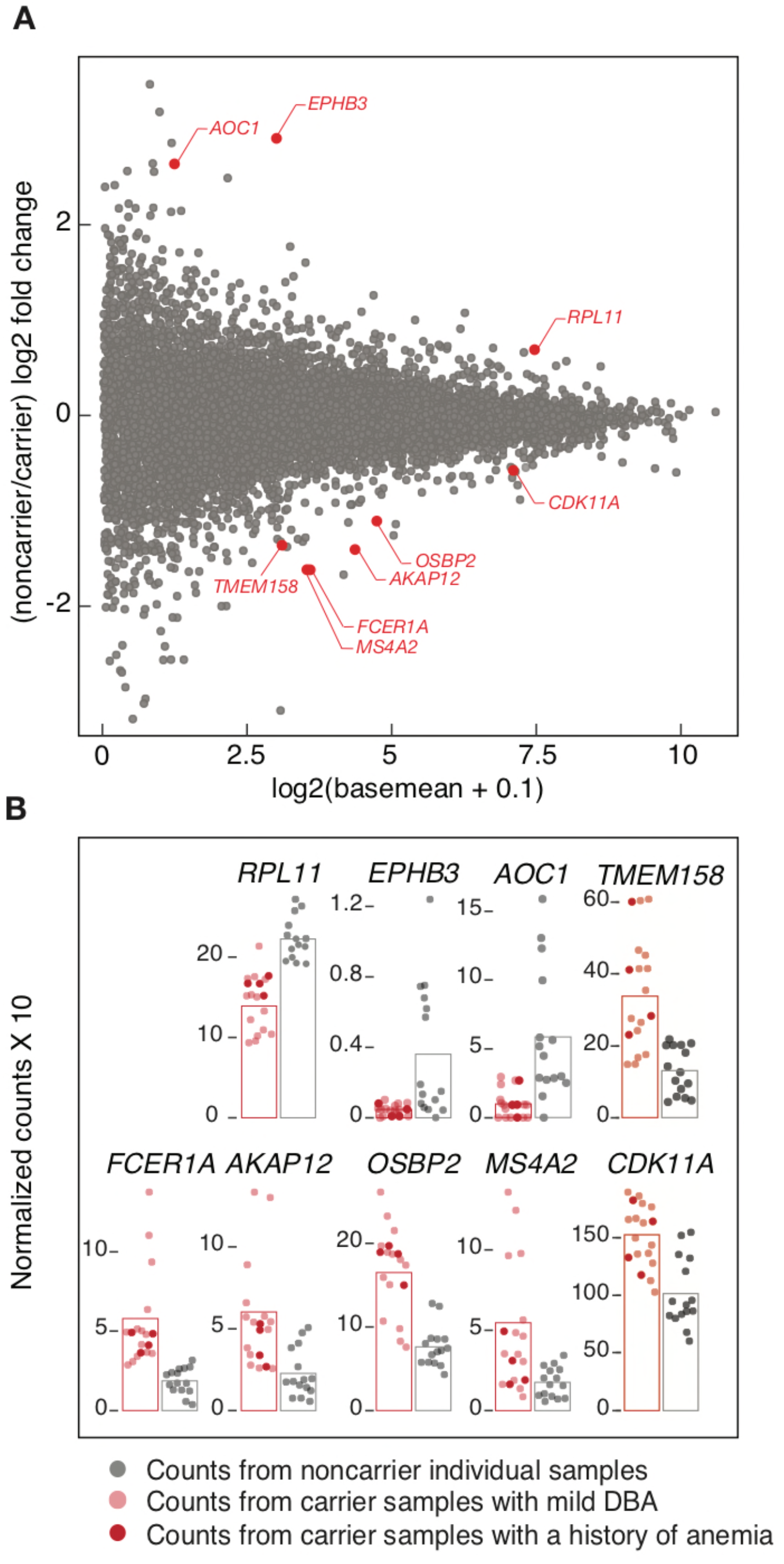
Shared gene expression changes among individuals that carry *RPL11* variant c.396+3A>G. Gene expression log2 fold changes (y-axis) and log2 base mean values per gene as calculated by DESeq2 were plotted. Y-axis represents log2 fold change of noncarriers compared to carriers. Red points are genes that are differentially expressed in two complementary approaches. (see Methods; Limma FDR cutoff <0.05, DESeq2 FDR cutoff <0.01). **B** Gene expression values per sample were plotted for differentially expressed genes in panel A. Noncarriers were labeled with grey. Carriers were shown in red and samples from individuals with a history of anemia are indicated with darker shade of red. The bars represent mean expression among all carriers (light red) and among noncarriers(grey).

Among the genes overexpressed in carriers **(Supplemental Table-3)**, we identified three implicated in cancer biology: *TMEM158, CDK11A* and *AKAP12. TMEM158* is a Ras induced senescence protein which is involved in proliferation arrest and its overexpression is linked to ovarian cancer [17]. Cyclin dependent kinase 11a (*CDK11A*), is a serine/threonine-protein kinase that controls cell cycle [18], and *AKAP12* is a tumor suppressor, inhibitor of cell migration and regulates cell-cycle progression through Src-mediated oncogenic signaling and cytoskeletal pathways [19,20]. An individual in this family with mild DBA symptoms was diagnosed with breast cancer at age 36, and there is an increased observed incidence of neoplasia including breast in the DBA registry. Thus, the observed significant overexpression of tumor suppressor genes; *TMEM158, CDK11A* and *AKAP12*, could be valuable understanding manifestation of increased incidence of neoplasia in DBA patients [21].

Among the under expressed genes in carriers, we identified *EPHB3*, an ephrin receptor that is expressed in early embryos and important for palate formation [22,23]. Among two family members with a history of anemia, cleft palate is observed in one, and congenital abnormalities are observed in the other. The observed differences in *EPHB3* transcript may relate to these symptoms.

Taken together, our findings suggest that individuals with the *RPL11* variant express the canonical isoform at a reduced expression level. We estimate that the c.396+3A>G variant produces ∼14% of the major splice isoform compared to the wild-type allele. Furthermore, this reduction in *RPL11* transcript leads to consistent but modest changes in RNA expression in a small number of genes that are involved in immunity, cell cycle, and cellular migration.

### Reduced expression of all ribosomal protein genes in carriers with mild DBA symptoms mirror the lower abundance of *RPL11*

Expression of ribosomal components is stochiometric [24,25]. Loss of expression coordination leads to unproductive energy consumption, proteotoxic stress [26], impaired cell proliferation, tumorigenesis and metastasis [examples: RPS3A [27], RPL5 [28], RPL15 [29–32].

Hence, we hypothesized that other ribosomal proteins may respond in a coordinated manner to reduced *RPL11* transcript expression. While individual ribosomal proteins were not identified as differentially expressed, we reasoned that a feedback regulation may lead to a coordinated regulation of the ribosomal subunits. Indeed, we observed that the expression of ribosomal protein genes was reduced modestly but consistently for both the small and large ribosomal subunit proteins in carrier individuals with mild DBA symptoms. We plotted the moderated log2-fold change (DESeq2) of all ribosomal proteins onto the human ribosome structure (PDB ID: 4UG0 [33]) to investigate any structural patterns to the feedback regulation (**Figure-4A**). We observed widely distributed reduced expression suggesting shared regulation among large and small ribosomal proteins with few exceptions (*RPL27A, RPL28, RPL39* and *RPL26, RPS29, RPS17*, and *RPS26;* example plots are shown in **Figure-4B**). This observation reveals a coordinated feedback regulation among other ribosomal protein genes in carrier individuals with mild DBA to compensate for the lower levels of *RPL11*.

**Figure-4:**
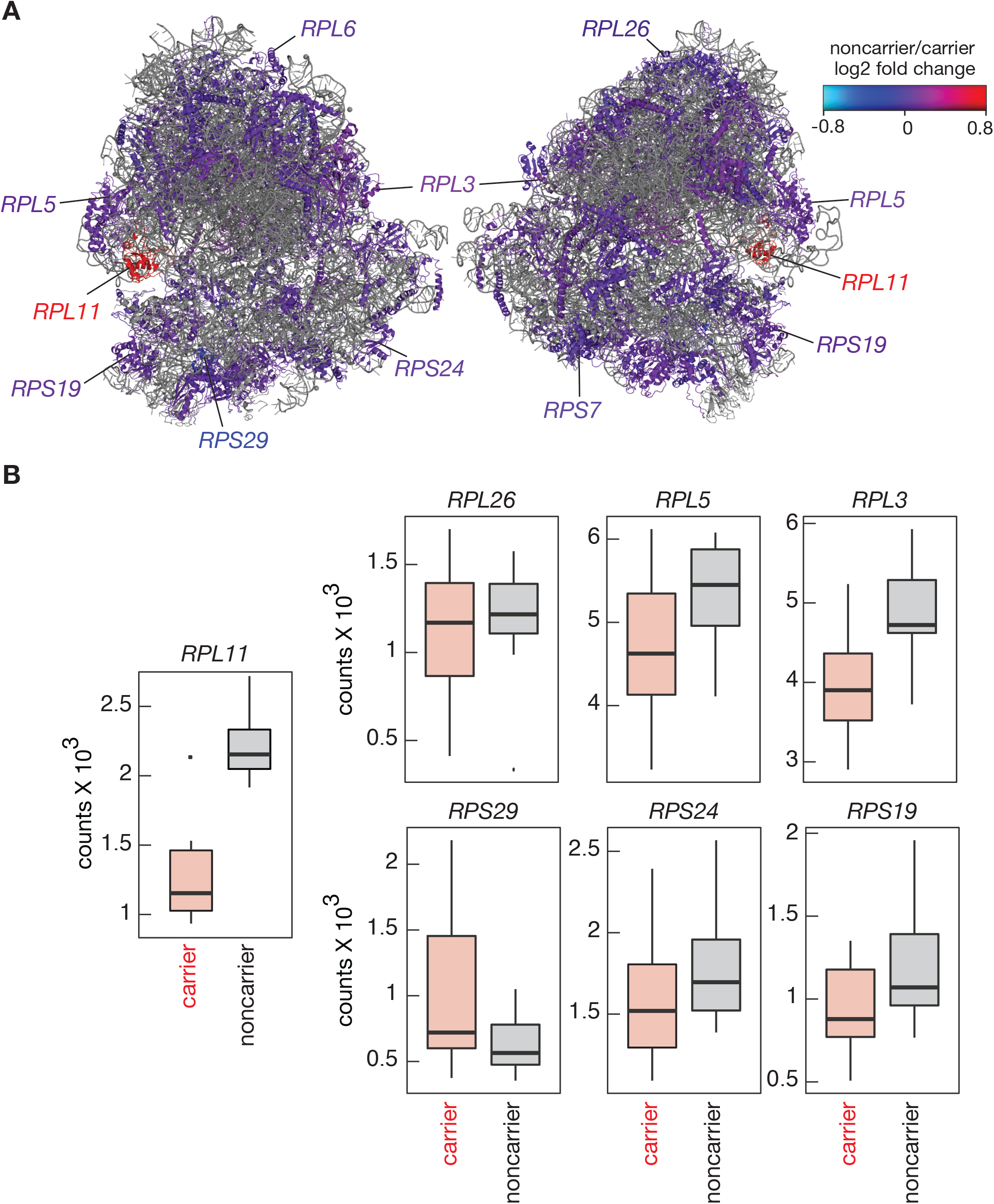
Effect of *RPL11* variant c.396+3A>G on the expression of both large and small ribosomal subunits. **A** Log2 shrinkage values (DESeq2) for each ribosomal protein gene were plotted in color scale on the human ribosome structure (PDB ID: 4UG0). **B** Distribution of normalized *RPL11* expression in carriers (light red) and noncarriers (grey) were plotted in the left panel. Right six panels display the distribution of three exemplary large (top) and three small (bottom) subunit ribosomal protein RNAs in carrier (red) and noncarrier (grey) individuals.

### Small and large ribosomal protein genes are significantly overexpressed in individuals with a history of anemia compared to carriers with mild or no symptoms

Given that carriers with mild DBA symptoms (one individual with no symptoms) have lower ribosomal protein gene expression compared to noncarriers (**Figure-4**), we expected to observe a similar trend of lower expression of ribosomal protein gene expression in carriers regardless of symptom severity. Surprisingly, we observed a different trend for the carrier individuals with a history of anemia: Both small and large ribosomal proteins are significantly overexpressed in these individuals compared to carrier individuals with mild symptoms (**Figure-5A and B**, ROAST, p-value=0.04; see Methods). Specifically, all ribosomal protein genes except four (*RPL11, RPS29, RPS17*, and *RPS26*) are overexpressed in carriers with a history of anemia (**Figure-5A)**. These results suggest that the compensation at the gene expression level for other ribosomal proteins in response to lower levels of *RPL11* transcript might be different in individuals with prominent symptoms compared to carriers with mild DBA.

**Figure-5:**
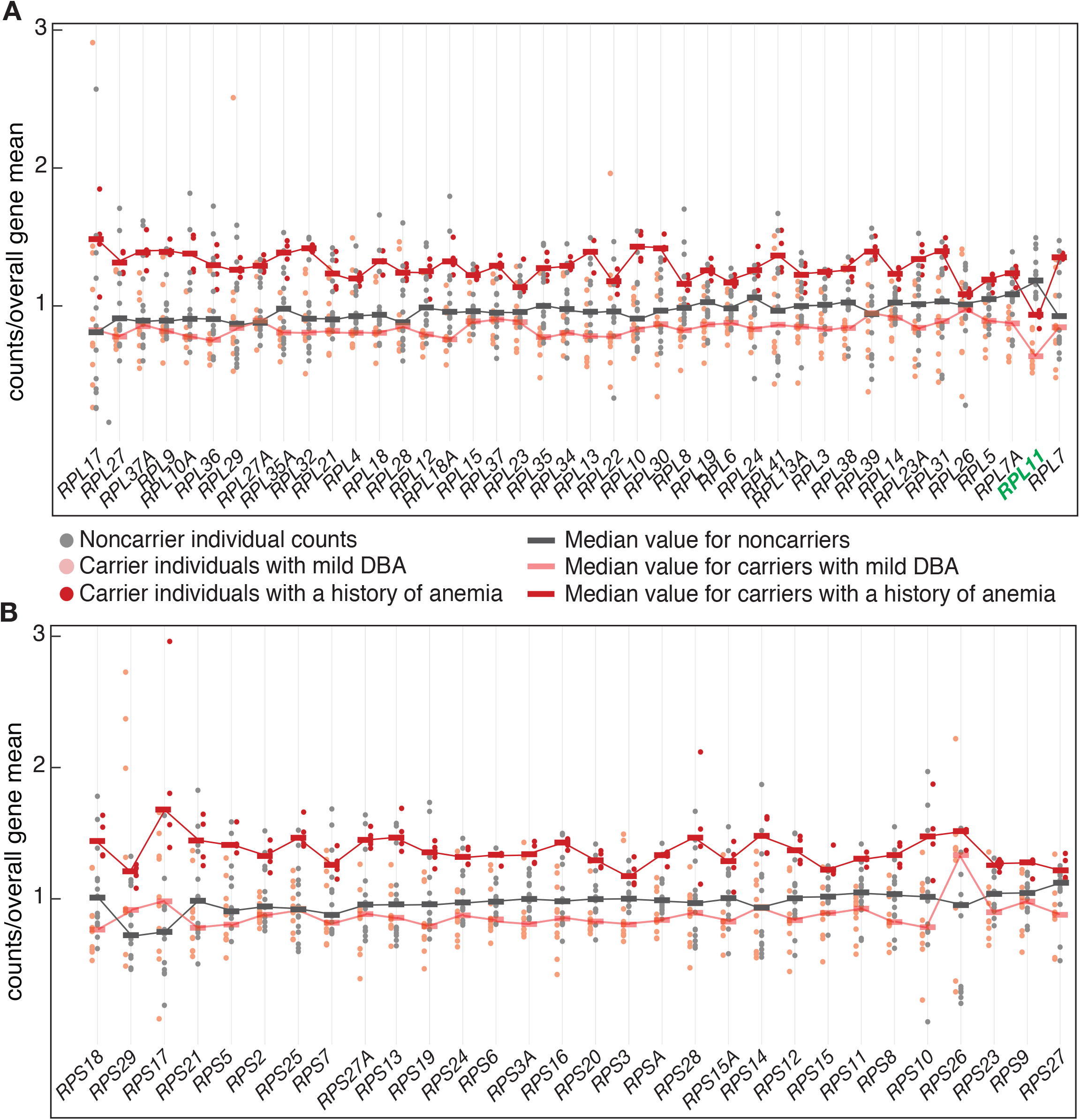
Large and subunit ribosomal protein mRNAs are differentially overexpressed in carrier individuals with a history of anemia compared to other carriers. Read counts for all ribosomal protein genes in each sample were divided to the mean of the cohort and were plotted for noncarriers (grey), and carriers. Carrier individuals with (dark red) or without (light red) a history of anemia were separated. Read counts that map to each large (top plot) and small (bottom plot) ribosomal protein transcript were divided by the mean for the corresponding ribosomal protein gene (y-axis). The thick color bars on the plot represent the median value for each gene, these bars are connected with a fine line to represent the overall pattern for each group. Large and small subunit ribosomal protein mRNAs except *RPL11* (highlighted in green) are significantly overexpressed in carriers with a history of anemia compared to carriers with mild DNA (ROAST–rotating gene set analysis, p value= 0.04). The difference between carriers without a history of anemia and noncarriers is not significant for ribosomal protein mRNA expression (ROAST, p value=0.1)

### Coordination between expression of mitochondrial components and *RPL11* are lost in all carriers

As previously implicated and shown in this study, feedback loops connect the expression of functionally related genes such as ribosomal proteins [24,25]. A novel hypothesis regarding variable expressivity of a phenotype is loss of gene expression coordination. Specifically, we aimed to identify clusters of genes that are expressed proportionally to *RPL11* yet lose this co-expression pattern once one copy of RPL11 is altered. This type of relationship cannot be detected through conventional differential expression analyses as *RPL11* loss would be predicted to lead to increased variance in the expression of the related genes and not their simple down- or over-expression.

To investigate this hypothesis, we used the variance of log-ratios of read counts between a given pair of genes as a function of *RPL11* carrier status. This metric, called emergent proportionality (‘theta E value’), enables discovery of gene pairs that are coordinately expressed if the ratio of their respective read counts is maintained among individuals regardless of their absolute expression level and carrier status [34,35]. Proportionality analyses formalize compositional data analysis methods for RNA expression and robustly infer underlying biological variables [35,36].

We identified 357 genes with higher variance of expression ratios compared to *RPL11* between carrier and noncarrier individuals (theta E < 0.15; **Supplemental Figure-6; Methods)**. Remarkably, carrier individuals dominated the variance for every single one of these top 357 genes (p < 2.2e-16; binomial test), suggesting an overall loss of coordination of gene expression in relation to *RPL11* in carriers **(Figure-6A)**. As a control, we conducted the same analysis with CDK11, a gene we found to be differentially increased in carriers (**Figure-3A)**. No significant results for directionality were obtained revealing that our results are not a simple consequence of differential expression **(Figure-6A)**.

**Figure-6:**
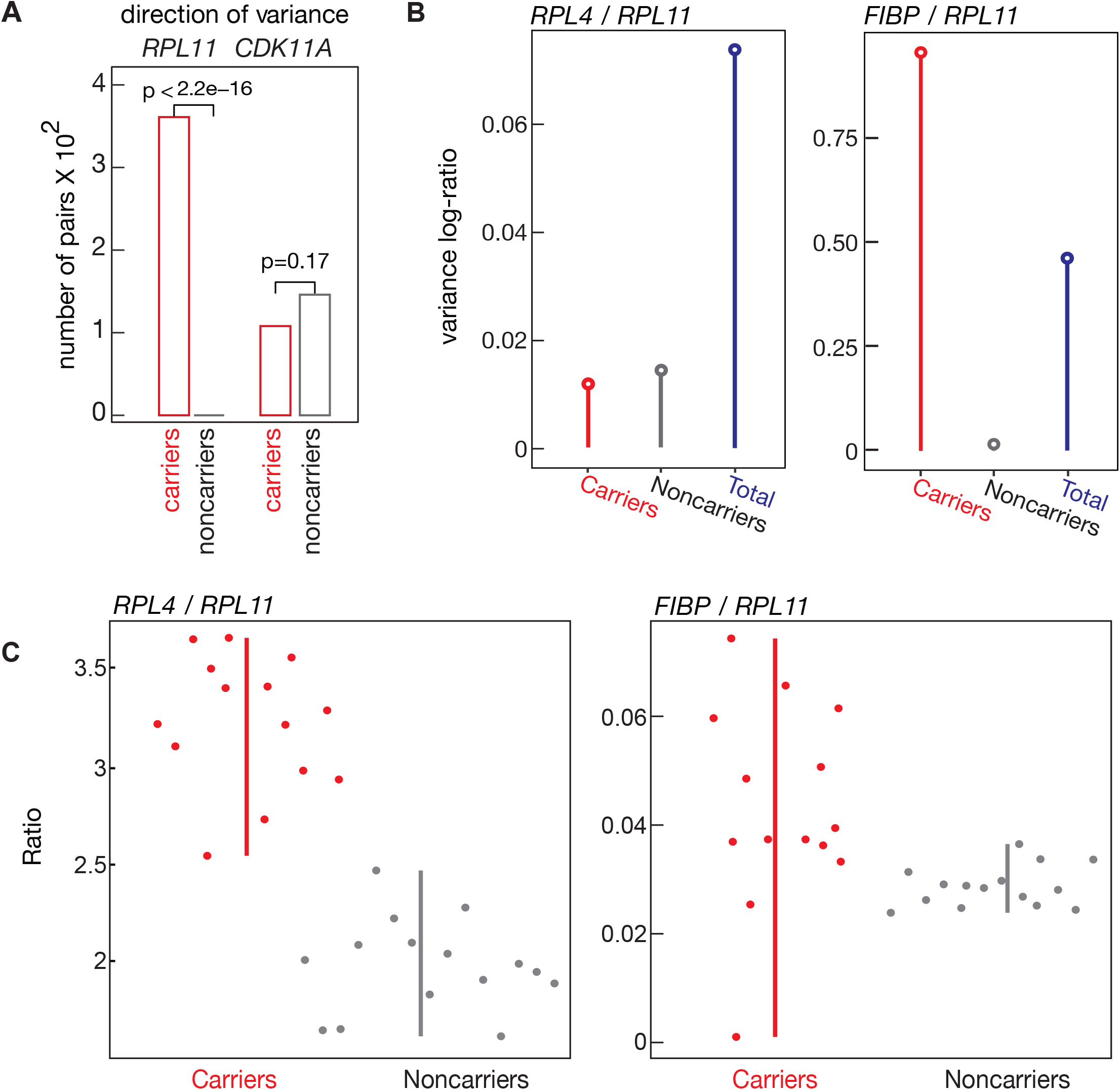
Emergent proportionality highlights loss of coordination between gene pairs. **A** The log-ratio of read counts of *RPL11* or *CDK11A* compared to all other genes were calculated. Variance of the log-ratios were used to calculate emergent proportionality (theta E) comparing carriers to non-carriers (Erb et al., 2017). For each gene pair with theta E <0.15–for proportionality to *RPL11–* or with theta E<0.25–for proportionality to *CDK11A*–, we determined the group (carrier or noncarriers) that dominates the total group variance. The significance of the bias towards either group was determined using a binomial test. **B** The total variance log-ratio (VLR) is plotted along with the VLR for noncarriers and carriers. Left panel: No genotype group dominates the total VLR, therefore *RPL4* and *RPL11* have a high theta E value (0.91). Right panel: VLR is higher in carrier samples for *FIBP / RPL11* (theta E = 0.01). **C** The ratio of two gene pairs for a sample is plotted and colored by *RPL11* status. These ratios are ordered by their sample index. Left panel: The ratio of *RPL4* and *RPL11* has similar variance by genotype, although the mean of the ratios differs. Right panel: For the ratio of *FIBP* and *RPL11*, carrier samples are spread out while noncarriers are distinctly more clustered

To better illustrate these expression patterns, we plotted one gene with similar and another with different variance of log-ratios (**Figure-6B, C**). *RPL4 to RPL11* (theta E value=0.91) ratio among carriers and noncarriers had similar variance suggesting that their expression coordination was similar in both groups (**Figure-6B, C**; left panels). On the other hand, *FIBP to RPL11* (theta E value=0.01) ratios were much more variable in carriers (**Figure-6B, C**; right panels).

We next determined the functional categories of genes that lost expression coordination with respect to RPL11. We found a very strong enrichment for genes related to mitochondria components and function **(Figure-7A and B**; **Supplementary Figure-7)**. In particular, there was a 45.4-fold enrichment for genes related to mitochondrial large ribosomal subunit, and a 10.6-fold enrichment for genes associated with the mitochondrial membrane (**Supplementary Table-5;** adjusted p-value < 0.05). We validated the robustness of these results to potential outliers via sub-sampling to calculate false discovery rates (**Supplementary Figure-6B**). Overall, these results reveal a loss of coordination between *RPL11* and mitochondrial components among carriers compared to noncarriers.

**Figure-7:**
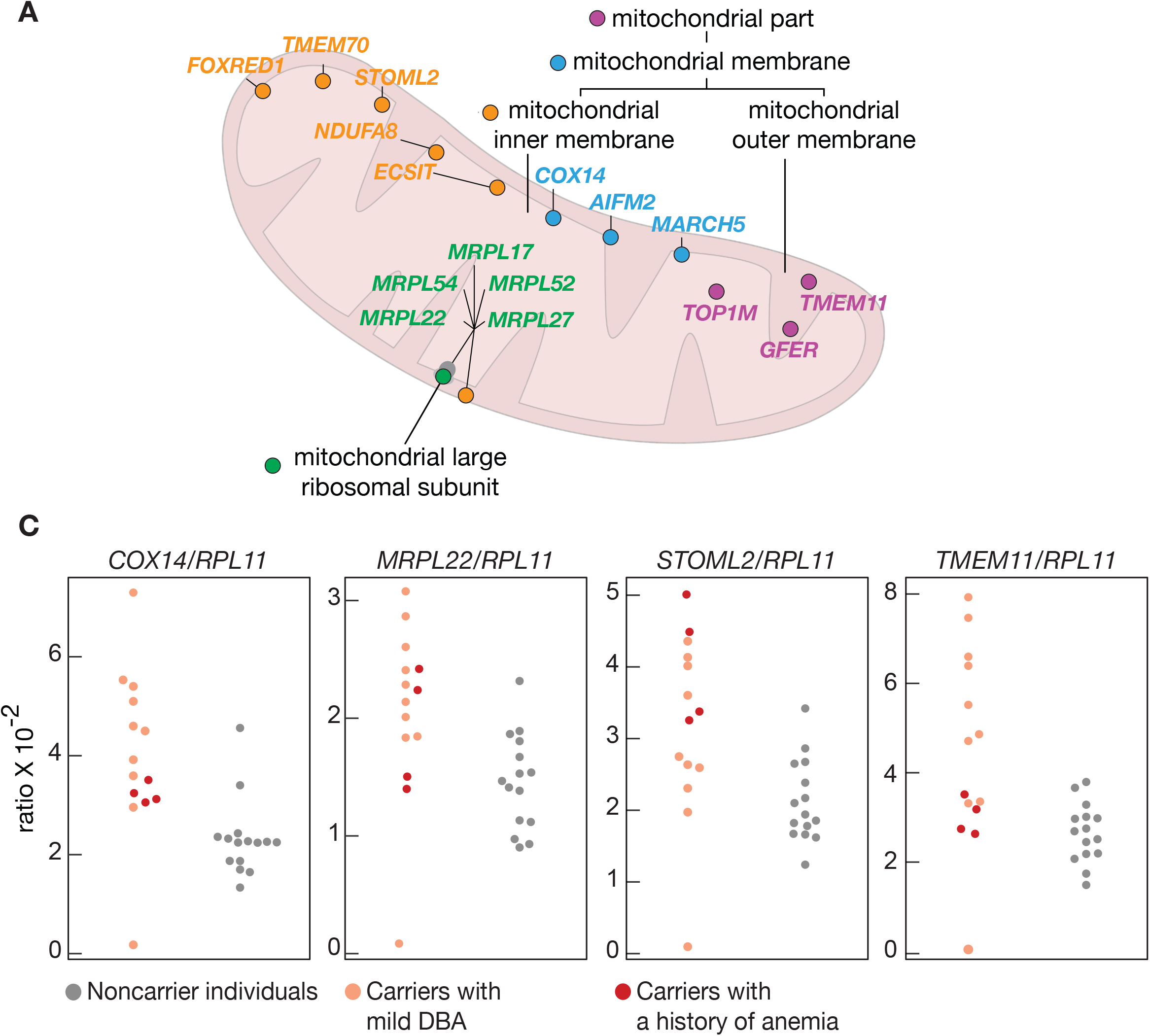
Expression of mitochondrial genes becomes uncoordinated with *RPL11* expression in carriers compared to noncarriers. **A** Genes with the lowest theta E values were highly enriched for mitochondrial localization as shown in the diagram [GO category significant enrichment analysis, adjusted p-value< 0.05, FuncAssociate 3.0 [58], Supplementary Table-5]. The only GO category that is not shown in this diagram belongs to cytoplasmic large ribosomal subunit components. **B** For selected mitochondrial genes, the ratio of read counts were plotted for the two groups.

## DISCUSSION

Variants in splicing regulatory sequences are major causative factors in a wide range of diseases including DBA [5,37]. Variants causing improper splicing can alter the encoded protein sequence and thus be detrimental to protein function. Here, we carried out an RNA sequencing study of a family spanning four generations with an inherited intronic variant affecting one of the two copies of *RPL11*. This noncanonical splice variant (c.396+3A>G) is causative for Diamond Blackfan Anemia (DBA) in this family with incomplete penetrance and variable expressivity.

We first determined the frequencies of aberrant splicing patterns attributable to this intronic variant. A previously missed splice junction with a 5’ splice donor site at +2 position relative to the canonical *RPL11* isoform has the highest abundance.

Interestingly, this isoform encodes a protein that is 75% identical to *RPL11* at its N-terminus and includes a late stop in the 3’UTR of the canonical isoform. Given the frameshifted polypeptide sequence in its C-terminal, this isoform may have proteotoxic consequences. In particular, SYO1, the dedicated chaperone for assembling RPL5 and RPL11 into 5S RNP complex, binds to the N-terminal of *RPL11* [38]. Thus, the aberrant RPL11 isoform may sequester SYO1 and abrogate ribosome biogenesis.

The expression of *RPL11* in carrier individuals is ∼57% of noncarrier individuals. If we assume that 50% of *RPL11* mRNA is transcribed from the variant *RPL11* locus, then we can deduce that ∼43% *RPL11* transcripts originating from the aberrantly spliced variant are degraded. This leaves only ∼7%–or lower if there is dosage compensation– of *RPL11* isoforms originating from the variant allele among the pool of *RPL11* transcripts.

Importantly, all carriers express the same set of aberrant splice isoforms with similar relative abundance. Therefore, symptom severity in this family is not attributable to difference in the splicing pattern among the carriers.

Among splice isoforms encoded by the *RPL11* variant, we were surprised that the second most abundant splice variant was anti-sense to the *RPL11* gene. These antisense junction reads are not present in any of the noncarriers, but surprisingly all of the carriers express them. Antisense exon-exon junctions have not been studied extensively in humans but were previously reported in *Plasmodium falciparum* [39] where they were detected in many transcripts. One hypothesis is that the antisense exon-exon junction reads result from splicing of an antisense transcript. The antisense RNA isoform we identified is only ∼80-100 nucleotides long, and we did not detect any other antisense read that maps to elsewhere in the *RPL11* coding region suggesting that antisense transcription of the *RPL11* gene would have to be highly unstable. We think that antisense transcription spanning only the exon4-exon5 region of the *RPL11* gene remains a more plausible alternative. Antisense oligonucleotides can be used as a tool to skip an exon or restore a cryptic splicing event [40]. However, the function of the detected antisense exon-exon junction remains unknown.

The family carrying *RPL11* variant fortunately display low penetrance for DBA symptoms (25%) compared to other DBA patients (∼75%). The two most affected patients have anemia in addition to congenital defects. The low penetrance might be specific to the ribosomal protein gene *RPL11*. Alternatively, the nature of the variant itself may be responsible. However, we observed highly similar splicing patterns for in all carrier individuals suggesting that any cis-acting splicing effects are unlikely to be explanatory.

To understand causes of incomplete penetrance and variable expressivity of DBA in this family, we investigated differential gene expression patterns in carriers with mild symptoms vs those with a history of anemia. Interestingly, carrier individuals with no history of anemia modestly downregulate other large and small ribosomal proteins suggesting a compensation mechanism for the lower expression of *RPL11*. On the other hand, carriers with a history of anemia significantly overexpress the majority of small and large ribosomal protein transcripts compared to other carriers. This could either be a response to symptom severity or directly be causative of the manifestations of DBA.

A lack of compensation in the expression of other ribosomal protein genes could lead to higher levels of proteotoxicity due to the increased abundance of unassembled ribosomal proteins. Majority of the overexpressed large subunit ribosomal protein mRNAs, if translated, would need to be degraded by the proteosome as they are usually unstable if not assembled into mature ribosomal subunits [41] Such toxicity hasn’t been documented in DBA patients before but would be an interesting avenue for further investigation.

Loss of coordination between ribosomal proteins and mitochondria may be yet another explanation for variable expressivity. When *RPL11* is reduced, expression of mitochondrial genes is not simply going up or down. Instead, mitochondrial genes are expressed with much higher variance compared to ribosomal protein genes. The increased variance in expression of mitochondrial components may lead to variable expressivity of DBA symptoms. For instance, the heme biosynthetic pathway which is important for red blood cell function originates in mitochondria [42]. Interestingly, patients with Pearson syndrome who harbor large mitochondrial lesions can frequently be misdiagnosed as having Diamond Blackfan anemia [43]. Furthermore, hematological abnormalities are observed in mitochondrial disorders [44]; and thus manifestation of DBA and variable expressivity could potentially be linked to the lack of coordination in mitochondrial components. Taken together, our study highlights the importance of unbiased systematic RNA expression analyses in families with non-canonical splice site variants that are causative for a Mendelian disorder.

## Acknowledgements

This work was supported in part by the Texas Rising Star Award [to E.S.C.] and National Institutes of Health [1R35GM138340-01 to E.S.C.; CA204522 to C.C.], Cancer Prevention and Research Institute of Texas [RR180042 to C.C.] and Welch Foundation [F-2027-20200401 to C.C.]. The authors acknowledge the Texas Advanced Computing Center (TACC) at The University of Texas at Austin for providing high performance computing and storage resources that have contributed to the research results reported within this paper. URL: http://www.tacc.utexas.edu. We also acknowledge the family for their participation in this research.

## MATERIALS AND METHODS

### Sample Collection and Human Subjects

Sample names, corresponding batches and index sequences for the RNA libraries are provided in **Supplementary Table-1**. We generated RNA-Seq libraries from at least two separate blood samples from 8 non-carriers, 5 carriers and 2 patients afflicted by anemia. In total, 43 RNA-Seq samples were generated and extensive quality control metrics were inspected prior to further analyses. A few of the RNAseq libraries were prepared from the same samples at different times to ensure we don’t observe bias due to batch effects. IRB approval as indicated in Carlston et al. 2017.

### RNA extraction and library preparation

RNA was extracted using Maxwell RSC instrument (Promega) from 50 ul of whole blood specimens with a white cell count greater than 50 000/µl. For depletion of ribosomal RNAs from the sample, 100 ng total RNA was mixed with 250 ng mixed pools of anti-sense rDNA oligos (**Supplemental Table-6**). RNA-DNA mix was incubated at 950C for 2 min, then the temperature was ramped down -0.10C per second to 450C. 1 ul Hybridase Thermostable RNase H (Lucigen, H39500) and 1 ul 10 X digestion buffer (500 mM Tris-HCl, 1M NaCl, 200 mM MgCl_2_) was preheated to 450C and added to the RNA-DNA mix. After RNAse H incubation for 1 hour at 450C, the samples were moved to 370C, and incubated for 1 hour in 50 µl DNAse reaction with 3 µl Turbo DNAse (ThermoFisher, AM2238) and 1X Turbo DNAse buffer. 300 µl solution containing 0.2 %SDS, 300 mM NaAcetate and 2 µl glycoblue was used to stop the reaction followed by acid-phenol extraction and ethanol precipitation. All of the precipitated RNA was converted into a sequencing library using SMARTer® smRNA-Seq Kit for Illumina (Thermo Fisher/Takara 635032). The kit protocol was used with the following specifics: RNA was initially hydrolysed for 4 minutes at 940C. The recommended additional AmpureXP cDNA purification was applied. The libraries were sequenced using 150 cycles on a HiSeq 2500. RNA libraries were prepared in four technical batches. Samples from individuals III.1, III.2, and III.4 were included in all of the batches to ensure the differences that are observed are not due to any batch biases. Sample names, corresponding batches and index sequences are provided in **Supplementary Table-1**.

### Preprocessing and Quality Control of RNA Sequencing Data

Sequencing reads were first trimmed to remove adapter sequences and low quality bases using cutadapt [45]. Specifically, we used the following parameters: “-a GATCGGAAGAGCACACGTCTGAACTCCAGTCACNNNNNNATCTCGTATGCCGTCTTCTGCTTG -A GATCGGAAGAGCACACGTCTGAACTCCAGTCACNNNNNNATCTCGTATGCCGTCTTCTGCTTG -A CAGATCGGAAGAGCGTCGTGTAGGGAAAGAGTGTAGATCTCGGTGGTC -u 6 -U 6 -q 22 -m 22”. Individual clipped fastq files were analyzed for quality control statistics using fastqc (https://www.bioinformatics.babraham.ac.uk/projects/fastqc/)Reads were aligned to the human genome using hisat2 [46] with default parameters. Alignment files were sorted and indexed using samtools [47]. Reads that map to protein-coding genes were counted using Rsubread’s featureCounts function for pair-end sequencing [48].

Transcript annotations were retrieved from Gencode’s v31 basic gene annotation file (06.2019) [49] and were filtered to include only genes with the \protein coding tag. Ensembl ids were converted to HGNC symbols through biomaRt’s hsapiens_gene_ensembl database and the getBM function with the following parameters: “attributes = c(“ensembl_gene_id”, “hgnc_symbol”), filter = “ensembl_gene_id”, values = ids, mart = ensemble_database)”.

### Quality Control Measures For RNA-Seq Data

The number of reads mapping to each protein-coding gene was compared for all pairwise combinations of samples. Samples were removed from further analyses if their mean Spearman correlation below 0.90 when compared to all others. **(Supplemental Figure-2)**. This filter resulted in the elimination of five samples. Next, we compared the annotated gender for each sample with that inferred from the data. Specifically, we profiled the gender of each individual based on their expression of chrY-linked genes and matched to the annotated gender. Our results indicated one sample displayed an apparent gender discrepancy compared to the annotation, as well as the inferred gender of other samples belonging to the same individual. Hence, this sample was removed. For the remaining samples, the distribution of counts was plotted, and no systematic bias as a function phenotype was observed **(Supplemental Figure-4A)**. Counts were transformed with DESeq2’s vst function and examined using principal component analysis (PCA) to cluster samples **(Supplemental Figure-4B)** (Love, Huber, & Anders, 2014). PCA analyses indicated that batch variables did not correlate with the first 4 principle components suggesting limited confounding technical variability to due batch effects **(Supplemental Figure-4C)**.

### Splicing Patterns in *RPL11*

For splicing pattern analysis of *RPL11*, samtools-1.9 was used to extract all sequencing reads that align to the *RPL11* gene locus (chr1:23,689,806-23,698,835) with MAPQ greater than 10. Integrative Genomics Viewer v2.6.0 (IGV) was used for visualization. All BAM files were converted to junction BED files in IGV by using the “export features’’ function while the entire *RPL11* gene is shown in IGV, to better quantify junction reads. To select junctions spanning exon 4 and 5 from the junction BED files, junctions starting between 23693000-23694999 and ending between 23695000-23695999 were selected. This strategy captures all junction reads that arise due to skipping exon 4. Because the start and end positions in the junction BED files are adjusted by the max read length on the junction, max read length on the start positions were added to the start positions and max read length on the end positions were subtracted from the end positions. The corrected junction start and end positions, read count, and the strand of junction were aggregated into a table in R version 3.6.3 **(Supplemental Table-2**). SpliceAIv1.2.1 was used to reconfirm the main effect of the variant in *RPL11* [50]. The effect of the most abundant splicing variant on translation was visualized in ExPASy (https:/www.expasy.org/).

### Antisense Junction Read Analyses

Genes transcribed from two strands were split into two gtf files using gencode’s v31 basic gene annotation (06.2019) file [49]. bedtools v2.27.1 was used to intersect sequencing reads with genes in the strand-separated gtf files, generating two files for each sample, one for genes transcribed from each strand. Samtools-1.9 was used to filter for reads with mapping quality (MAPQ) greater than 10. awk was used to select antisense junction reads from the alignment files. R version 3.6.3 was used to quantify each antisense junction read and filter for junctions that had more than 3 reads in at least 5 samples. The selected junction reads were visualized in Integrative Genomics Viewer v2.6.0.

### Splice Region Variants in the Human Genome

Splice variant data from gnomAD v3 was used for analyzing splice region variants. Loss-of-function (LoF) metrics data in gnomAD v2.1.1 [51] was used to filter genes by probability of loss-of-function intolerance (pLI), with thresholds of pLI greater than 0.96 or less than 0.04. The value 0.96 was taken from the pLI of *RPL11*. Python 3.7.4 was used to download and aggregate the data for ribosomal protein genes and genes with matching pLI from the last step, from the gnomAD API. The table was read into R version 3.6.3 and analyzed.

### Differential Expression Analyses

Differential expression was conducted with the R packages DESeq2 and limma-dream. Carriers, including DBA patients were compared to noncarriers, with the only covariate for the model being *RPL11* variant status. For DESeq2, technical replicates were collapsed by summing the counts of that individual’s technical replicates. Differentially expressed genes were identified using the standard pipeline in [52]. Genes with an adjusted p-value of less than equal to 0.01 were deemed significant [53]. As a complementary approach, Dream was utilized in conjunction with limma to accommodate the repeated measures in experimental design through a linear mixed-effects model [54–56]. The counts of technical replicates were summed for each sample. Independent samples of an individual were then modeled with a linear mixed-effects model along with *RPL11* status [56]. The results for both analyses were combined by considering genes with a p-value less than 0.05 in limma-dream and an adjusted p-value of less than 0.01 in DESeq2 as potentially differentially expressed between groups. Rotation gene set testing for linear models was employed through the FRY function in limma to test group differences in ribosomal genes [57]. Comparisons with p-values under 0.05 were deemed significant.

### Differential Proportionality

For differential proportionality, the definition established for emergent proportionality by Erb *et al*. 2017, was used [34,35]. Modifications to the methods regard robust sampling, which randomly samples an equal number of individuals to represent each group delimited by *RPL11* status for 100 permutations. For each permutation, theta E, a measure of emergent proportionality, was calculated between *RPL11* and all other genes, using *RPL11* status as the biological variable of interest. The average of these permutations was utilized in further analysis **(Supplemental-Figure-6A)**. The false discovery rate was estimated by randomly assigning *RPL11* status to all individuals, calculating theta E with the modified methods described above, repeating for 100 permutations, and comparing expected to observed values **(Supplemental-Figure-6B)**. For the ratio of each gene to *RPL11*, one group will have a greater amount of variance than the other. The direction of variance is defined as which *RPL11* status possesses the dominating amount of variation in the ratio of two genes. With this classification of direction of variance, a binomial test was used to determine whether the direction of variance was significantly different between the group of interests with the assumption this direction is equally distributed. Theta E less than 0.15, representing the most significant results, was used to identify gene-pairs where a single group contributes the majority of the variance. Gene ontology analysis was performed through FuncAssociate 3.0 using an ordered list of genes (Theta E < 0.15), and terms were kept with an adjusted p-value cutoff of 0.05 [58].

## Figure Legends

**Supplemental Figure-1: An exemplary of IGV snapshots of *RPL11* gene at exon4-intron4 region from example carrier and non-carrier individuals**. Few reads that map to *RPL11* variant c.396+3A>G was used as a confirmation of the carrier status.

**Supplemental Figure-2: Plot of Spearman correlation across RNA samples and individuals**. 5 samples with lower spearman correlation values (<0.9) were discarded from further RNA expression analysis. These 5 samples were further investigated in Figure S3. The mean correlation value across samples after the sample removal is 0.97.

**Supplemental Figure-3: Samples with <0.9 Spearman correlation from Figure S1 generally have lower percentage of genome mapping reads**. Samples II.8_s2, III.3_s1_t3, III.4_s1_t4, which were removed from further analysis, generally have low counts and percentages of mapped reads in comparison to the other example samples; II.6_s2 and III.1_s1_t3.

**Supplemental Figure-4: A** Normalized assigned counts were plotted for each sample. Grey, light red and dark red points represent noncarriers, carriers with mild DBA and carriers with a history of anemia, respectively. **B** PCA analysis plot is shown for each sample annotated with individual identifier from Figure 1, followed by a number that stands for sample number for the particular individual. RNA expression libraries were prepared in four batches and three samples were included in all of the batches. **C** Libraries that were prepared in each batch was colored accordingly (batch 1–purple, 2– blue, 3–green, 4–orange) in the two PCA plots; first two variance components are plotted left, third and fourth components are plotted on the right. Note that PCA plots in B and C are the same, where each sample were labeled with a sample name, or a batch number respectively.

**Supplemental Figure-5: Shared gene expression changes among carrier individuals with mild DBA compared to noncarriers. A** Gene expression log fold changes (y axis) and log base mean values per gene were plotted on the left large plot. Red labeled points are significant genes that are differentially expressed (Limma Analysis FDR cutoff<0.05, DESeq2 Analysis FDR cutoff<0.01). **B** Mean gene expression of carriers with mild DBA and a history of anemia (y axis on the left and right plots respectively) are compared to noncarriers (x axis). Red points on the plot represent significantly over and under expressed genes in carriers with mild DBA in comparison to noncarriers.

**Supplemental Figure-6:** Implementation of individual sampling allows for more robust emergent proportionality results. **A** Equal group sizes are formed by randomly sampling noncarriers individuals equal to the number of carrier individuals (n=7). One sample from each individual in the pool is then selected. Theta E is calculated with this reduced pool. This process is repeated for 100 permutations and the results are averaged. **B** False discovery rates (FDR) are calculated in a similar manner, where samples are first randomly mixed. The number of gene pairs with a theta E below cutoffs is then calculated.

**Supplemental Figure-7: The loss of coordination of mitochondrial component genes to *RPL11* in carriers are plotted for each gene**. The 16 mitochondrial related genes that are (theta E < 0.15) in emergent proportionality with respect to *RPL11* are plotted by their ratio to *RPL11* and are colored by phenotype.

**Supplemental Table-1:** Details about the individual samples and RNA libraries are provided.

**Supplemental Table-2:** Related to Figure 2. Raw number of reads per sample are provided for each of the splice junctions that are plotted in Figure-2.

**Supplemental Table-3:** Log fold changes, mean expression values as calculated by Dream and DESeq Analysis are provided with P values. First tab provides the list for comparison of noncarriers to all carriers. Second tab provides the list of comparison of noncarriers to carriers without a history of anemia only.

**Supplemental Table-4:** DESeq normalized reads per sample are provided. Please check family diagram in Figure 1 for the status of each individual.

**Supplemental Table-5:** Gene ontology (GO) analysis was performed for the genes with a theta E less than 0.15 with respect to *RPL11* using FuncAssociate 3.0 [58]. The list is provided for all significant GO terms with an adjusted p-value below 0.05.

**Supplemental Table-6:** Antisense ribosomal DNA oligos used for the rRNA depletion for RNA-seq library preparation.

