## Supplemental Figures for "Loss of coordinated expression between ribosomal and mitochondrial genes revealed by comprehensive characterization of a large family with a rare mendelian disorder"

### Supplemental Figure-1

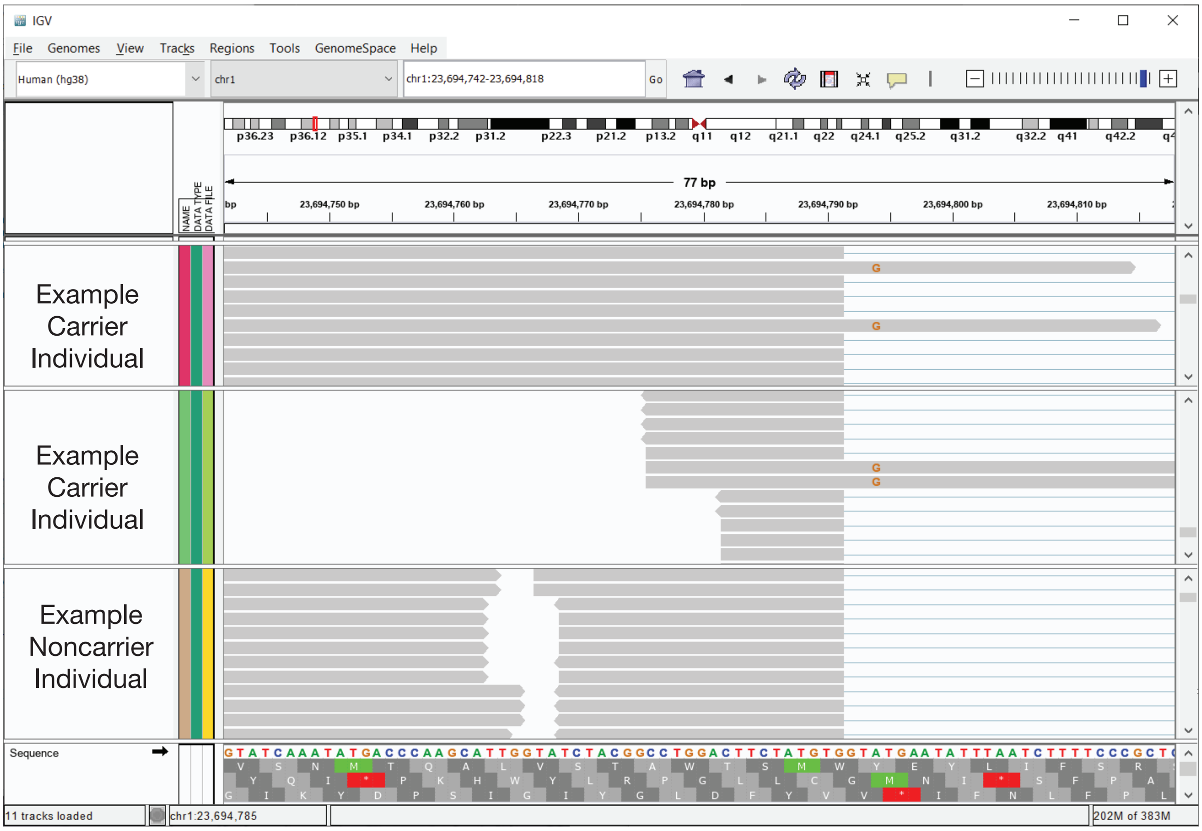


**Supplemental Figure-1: An exemplary of IGV snapshot of *RPL11* gene at exon4-intron4 region from example carrier and non-carrier individuals.** Few reads that map to *RPL11* variant c.396 3A>G was used as a confirmation of the carrier status.

#### Supplemental Figure-2

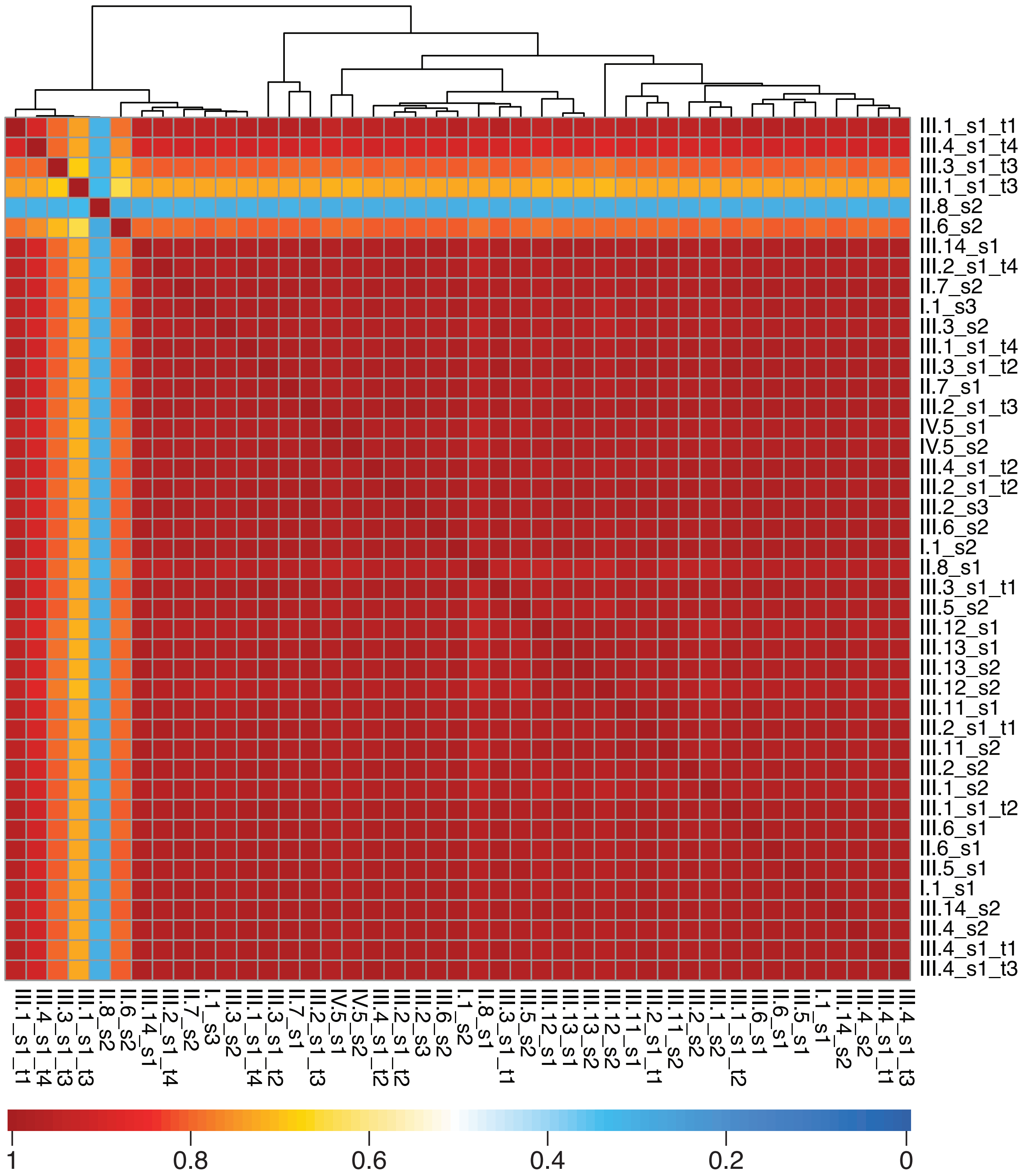


**Supplemental Figure-2: Plot of spearman correlation across RNA samples and individuals.** 5 samples with lower spearman correlation values (<0.9) were discarded from further RNA expression analysis. These 5 samples were further investigated in Figure S3. The mean correlation value across samples after the sample removal is 0.97. Across samples, example: III.4, refers to the individual; example: s1 refers to the biological sample taken, example: t1 refers to a technical replicate (RNAseq library was prepared from the same sample at different times). The libraries were prepared in four batches. A few of the samples were included and processed as technical replicates in all of the batches to ensure that a batch bias doesn't exist (Supplementary Table-1).

#### Supplemental Figure-3

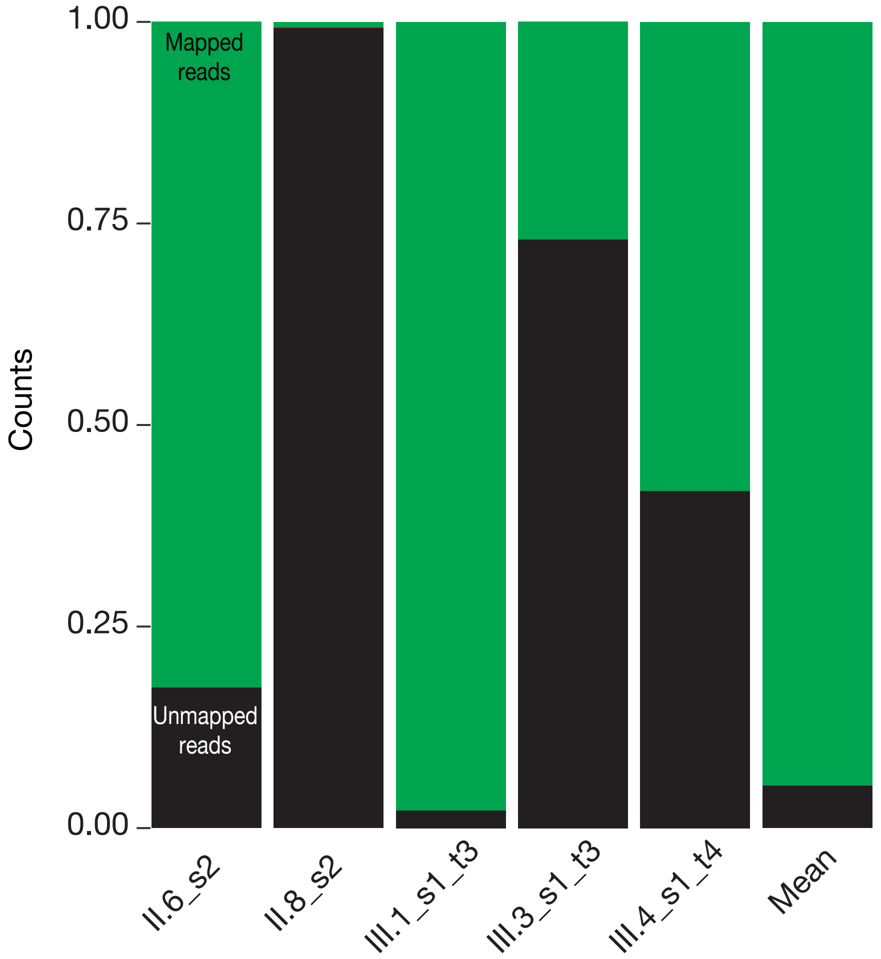


**Supplemental Figure-3: Samples with <0.9 Spearman correlation from Figure S1 generally have lower percentage of genome mapping reads.** Samples II.8_s2, III.3_s1_t3, III.4_s1_t4, which were removed from further analysis, generally have low counts and percentages of mapped reads in comparison to the other example samples; II.6_s2 and III.1_s1_t3.

#### Supplemental Figure-4

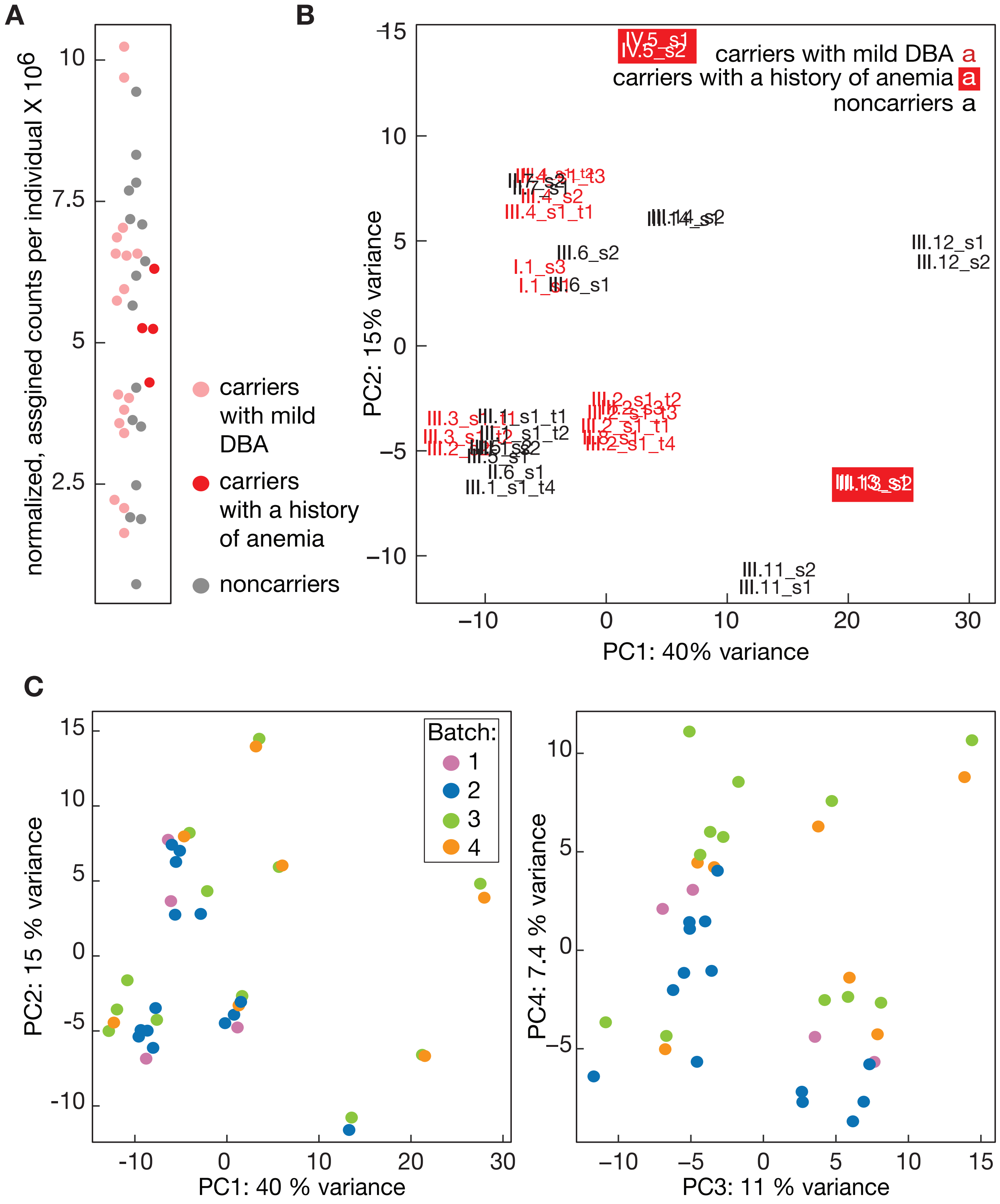


**Supplemental Figure-4: A** Normalized assigned counts were plotted for each sample. Grey, light red and dark red points represent noncarriers, carriers with mild DBA and carriers with a history of anemia, respectively **B** PCA analysis plot is shown for each sample annotated with individual identifier from Figure 1, followed by a number that stands for sample number for the particular individual. RNA expression libraries were prepared in four batches and three samples were included in all of the batches. **C** Libraries that were prepared in each batch was colored accordingly (batch 1–purple, 2–blue, 3–green, 4–orange) in the two PCA plots; first two variance components are plotted left, third and fourth components are plotted on the right. Note that PCA plots in B and C are the same, where each sample were labeled with a sample name, or a batch number respectively.

#### Supplemental Figure-5

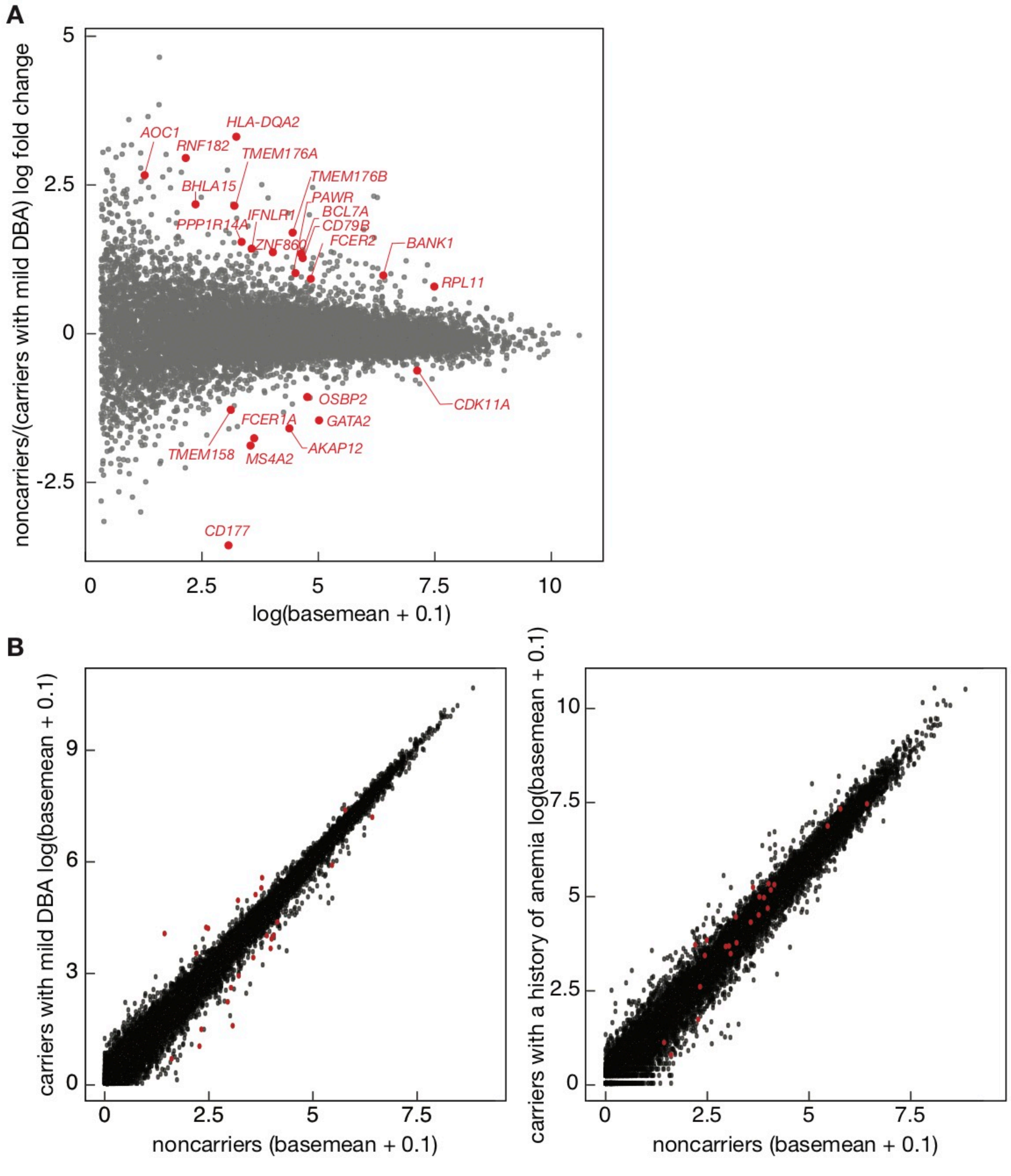


**Supplemental Figure-5: Shared gene expression changes among carrier individuals with mild DBA compared to noncarriers. A** Gene expression log fold changes (y axis) and log base mean values per gene were plotted on the left large plot. Red labeled points are significant genes that are differentially expressed (Limma Analysis FDR cutoff<0.05, DESeq2 Analysis FDR cutoff<0.01). **B** Mean gene expression of carriers with mild DBA and a history of anemia (y axis on the left and right plots respectively) are compared to noncarriers (x axis). Red points on the plot represent significantly over and under expressed genes in carriers with mild DBA in comparison to noncarriers.

#### Supplemental Figure-6

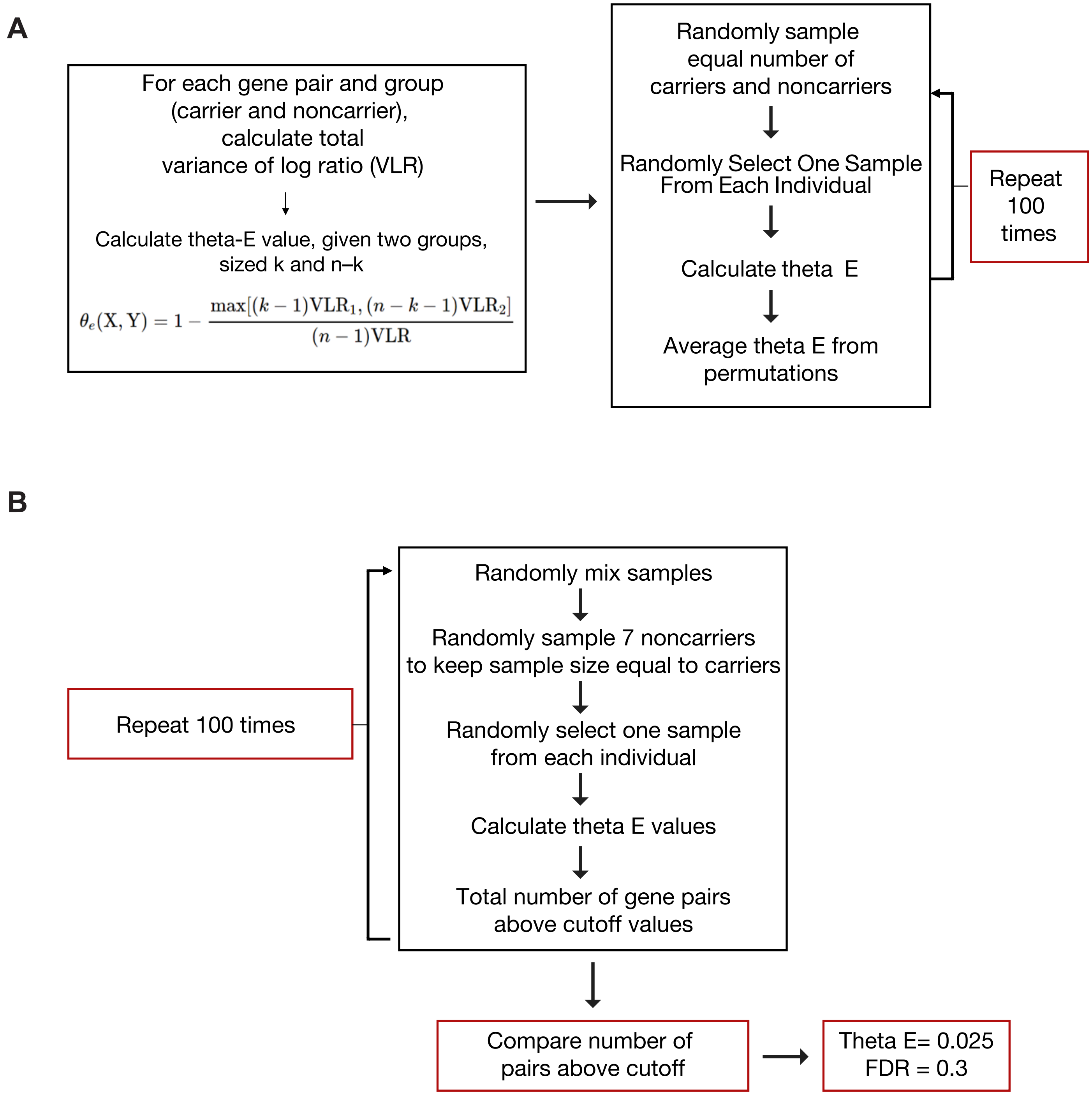


**Supplemental Figure-6: Implementation of individual sampling allows for more robust emergent proportionality results. A** Equal group sizes are formed by randomly sampling noncarriers individuals equal to the number of carrier individuals (n=7). One sample from each individual in the pool is then selected. Theta E is calculated with this reduced pool. This process is repeated for 100 permutations and the results are averaged. **B** False discovery rates (FDR) are calculated in a similar manner, where samples are first randomly mixed. The number of gene pairs with a theta E below cutoffs is then calculated.

.

#### Supplemental Figure-7

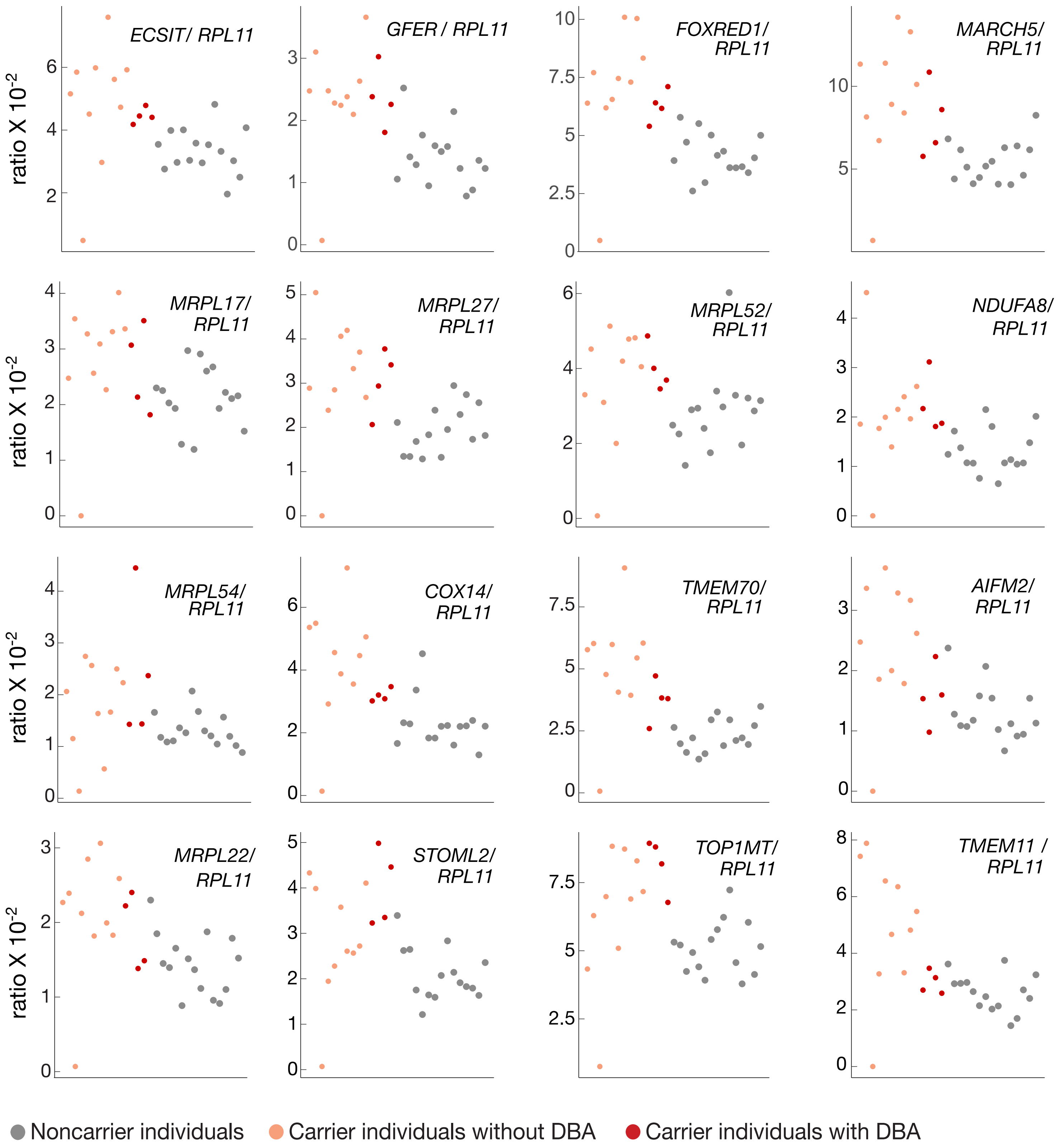


**Supplemental Figure-7: The loss of coordination of mitochondrial component genes to *RPL11* in carriers are plotted for each gene.** The 16 mitochondrial related genes that are (theta E < 0.15) in emergent proportionality with respect to *RPL11* are plotted by their ratio to *RPL11* and are colored by phenotype.
